# Fully Automated Sequential Immunofluorescence (seqIF) for Hyperplex Spatial Proteomics

**DOI:** 10.1101/2023.07.07.548135

**Authors:** François Rivest, Deniz Eroglu, Benjamin Pelz, Joanna Kowal, Alexander Kehren, Maria Giuseppina Procopio, Pino Bordignon, Emilie Pérès, Marco Ammann, Emmanuel Dorel, Sylvain Scalmazzi, Lorenzo Bruno, Matthieu Ruegg, Gabriel Campargue, Gilles Casqueiro, Lionel Arn, Jérôme Fischer, Saska Brajkovic, Pierre Joris, Marco Cassano, Diego Dupouy

**Affiliations:** Lunaphore Technologies SA, Tolochenaz, Switzerland

## Abstract

Tissues are complex environments where different cell types are in constant interaction with each other and with non-cellular components. Preserving the spatial context during proteomics analyses of tissue samples has become an important objective for different applications, one of the most important being the investigation of the tumor microenvironment. Here, we describe a multiplexed protein biomarker detection method on the COMET instrument, coined sequential ImmunoFluorescence (seqIF). The fully automated method uses successive applications of antibody incubation and elution, and in-situ imaging enabled by an integrated microscope and a microfluidic chip that provides optimized optical access to the sample. We show seqIF data on different sample types such as tumor and healthy tissue, including 40-plex on a single tissue section that is obtained in less than 24 hours, using off-the-shelf antibodies. We also present extensive characterization of the developed method, including elution efficiency, epitope stability, repeatability and reproducibility, signal uniformity, and dynamic range, in addition to marker and panel optimization strategies. The streamlined workflow using off-the-shelf antibodies, data quality enabling quantitative downstream analysis, and ease of reaching hyperplex levels make seqIF suitable for immune-oncology research and other disciplines requiring spatial analysis, paving the way for its adoption in clinical settings.

## Introduction

Recent advances in cancer immunotherapies and novel combination therapies have resulted in a surging interest in the study of the tumor microenvironment (TME) [1, 2]. The TME is a heterogeneous biologically functional region that consists of a variety of cell types including tumor infiltrating immune cells interacting with malignant cells, in addition to stromal cells and endothelial cells as well as non-cellular components such as extracellular matrix and cytokines. Hence, it is essential to preserve the spatial context to obtain the full extent of biological information while studying the complex cellular interactions within the TME, which in turn can provide unparalleled insights into predicting response to therapies. A well-known example of leveraging spatial information in the TME is the development of the Immunoscore, a method using the density of infiltrating CD3 and CD8-positive immune cells within the tumor and its invasive margin as a metric [3, 4]. This method demonstrated superior performance as prognostic indicator for colorectal cancer (CRC) in certain cases when compared to other metrics such as staging or microsatellite instability [5].

The analysis of the interaction between TME and the immunological components is mainly performed through the pathological evaluation of biological tissue specimens. Hematoxylin & Eosin staining (H&E) is the gold standard for obtaining information on tumor morphology, often used in differential diagnosis [6]. Immunohistochemistry (IHC) adds an additional dimension to the analysis of tissue specimens – the ability to visualize the expression of specific proteins while preserving the tissue architecture. IHC is a robust and simple technique, both in regards of assay development and visualization, but it has several limitations when applied for the analysis of the TME. Due to the nature of the chromogenic visualization, the standard IHC method is limited to one marker per slide. Alternative implementations can introduce multi-marker detection capability, which is, however, still limited to low-plex use cases [7]. IHC also lacks true quantitative capability, which requires coupling it with binary or quantized scoring methods [8].

Multiplex immunofluorescence (mIF) is a technique which overcomes these limitations, offering highly quantitative data with multi-marker capability, with the potential to scale to hyperplex levels [9–11]. One of the most popular mIF approaches involves the use of tyramide signal amplification (TSA) and multispectral imaging, where samples are subjected to successive incubations of primary and horseradish peroxidase (HRP)-conjugated secondary antibodies followed by HRP-mediated in-situ deposition of the tyramide-fluorophore, and antibody removal [12, 13]. A single imaging step at the end of the protocol captures the cumulative fluorescent signals. The multiplex capability of this approach is inherently limited by the crosstalk between the emission spectra of different fluorophore signals, resulting in assays typically limited to 6-plex. It can potentially be enhanced to up to 9-plex with specialized spectral unmixing methods [14]. The use of custom antibody conjugation techniques such as DNA- or metal-based barcoding can remove some of the barriers associated with mIF and enable increased multiplexity [15, 16] at the cost of adding assay complexity and limiting the spectrum of research questions that can be addressed due to the necessity of using custom antibodies. Moreover, the readout can use pseudo-images and result in permanent damage on the interrogated tissue section, which in turn, prevents downstream processing steps such as H&E staining [17, 18]. Cyclic IF approaches aim to increase multiplex capacity while sticking to standard antibodies, applying imaging and signal removal steps between antibody incubation cycles [19, 20]. The main limitations of these approaches are: (i) the excessive need for manual handling in repeated cycles, (ii) long turnaround times, and (iii) potential sample degradation due to either repeated sample mounting and unmounting or harsh conditions to bleach or remove the signal [21].

We introduce a straightforward sequential immunofluorescence (seqIF) methodology for the spatial detection of protein biomarkers on a wide range of tissue types, which consists of iterative sets of staining, imaging and the elution steps of macromolecular antibody complexes while preserving tissue antigenicity and morphology. We then describe the implementation of seqIF on COMET™, a fully automated platform based on a unique microfluidic technology, that works with off-the-shelf reagents. This platform has been shown to provide the “sample in - image out” capability for the detection of 40 markers in a single automated run, without tissue damage, eliminating the need for strenuous manual handling and offering reduced turnaround times, while still allowing the use of established and characterized (unconjugated) primary antibodies. The microfluidic technology behind COMET™ has first been demonstrated for IHC [22], with results showing highly reduced ambiguity in HER2 scoring. The precise immunofluorescent staining ensured by microfluidics technology, when coupled with image analysis methods, was shown to have high correlation with molecular expression patterns of biomarkers [23]. The first multi-marker fluorescent immunostaining results with this technology were shown in a TSA multiplexing setting [24], and seqIF assays developed with the predecessor of COMET™ enabled the spatial analysis of the TME [25]. The technology behind seqIF has also been used to generate insight into biological outcomes in different tissue types [26, 27]. Here, we explain in detail the established seqIF workflow, including marker optimization and panel development strategies, along with results from hyperplex experiments.

## Results and Discussion

### COMET and automated seqIF overview

An overview of the different aspects that enable the development of the automated seqIF protocol for hyperplex assays is demonstrated in Figure 1. Figure 1a shows the COMET™ instrument, which comprises a reagent module, an integrated microscope and multiple automated stainers. The reagent module contains 20 reservoirs to be used for primary antibodies, corresponding to walk-away automation for a 40-plex assay with a configuration of combining antibodies from 2 different species, isotypes or subclasses per each staining cycle. Larger volume reservoirs are used for buffers and detection system reagents such as secondary antibodies. The automated stainers are placed on a motorized stage to move the sample ready to be imaged under the integrated microscope while staining the following one in parallel, enabling higher throughput.

**Figure 1:**
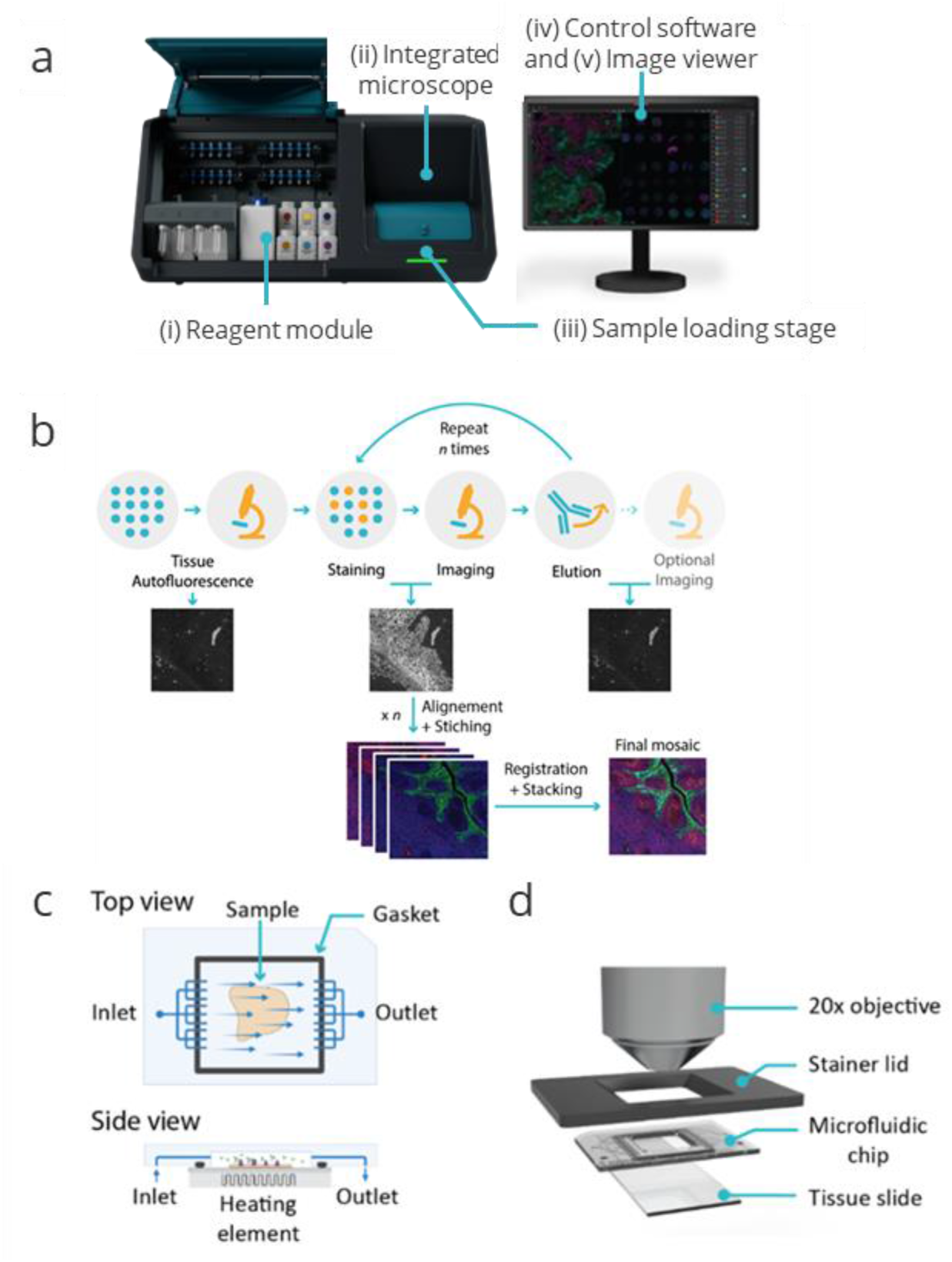
Overviews of COMET™ platform and seqIF method, enabling full automation and walk-away execution of multiplex immunofluorescence assays. **(a)** COMET™ platform overview. (i) A reagent module composed of 20 small volume reservoirs for reagents, typically used for primary antibodies, 4 mid-size reservoirs, typically used for detection reagents, and 7 large volume reservoirs for ancillary buffers; (ii) an integrated epifluorescence microscope acquiring automated mosaic scans of stained samples, (iii) a sample loading/unloading interface with a rotary stage that accommodates 4 automated stainers (iv) a control software for protocol preparation and execution; and (v) an imaging viewer for rapid visualization of multiplexed immunofluorescence results. **(b)** Sequential immunofluorescence method (seqIF). Tissue autofluorescence is automatically acquired as a baseline for downstream background removal. Samples are automatically stained and imaged for DAPI and 2 markers of interest. Elution - Signal removal by chemical removal of antibodies. Staining, imaging, and elution are repeated *n* times (*n:* number of multiplexed staining steps). Images are automatically stitched, aligned, and stacked on a final OME-TIFF file. **(c)** Top and side views of the microfluidic imaging chip consisting of inlet, outlet and a fluidic network that enable uniform exposure of reagents over the tissue sample. A gasket at the edges of the staining area ensures a hermetic sealing. A heating element underneath is used to control the temperature of each step in the seqIF protocol. **(d)** Integrated imaging assembly. The microfluidic chip contains an imaging window that allows the integration of in-situ fluorescence microscopy and permits the direct imaging of the sample.

Figure 1b shows an illustrative flowchart of the seqIF method on COMET™. It starts with a tissue autofluorescence acquisition step, where the output can be used in image post-processing if autofluorescence subtraction option is preferred. Afterwards, the iterative section of the protocol begins, which consists of short incubation steps with primary and fluorescently labeled secondary antibodies, followed by imaging and elution to gently strip off primary and secondary antibodies. At every cycle, two primary antibodies can be applied, followed by detection with two complementary fluorescently labeled secondary antibodies, each targeting one of primary antibody, and matching the excitation and emission spectra of TRITC and Cy5 channels available in the instrument. To further improve the turnaround time and throughput, the platform has the capability to execute staining and imaging cycles in parallel between different slides, reducing the overall protocol time by half when multiple samples are run in a single protocol. After each elution step, an optional imaging step can also be performed. All images registered during the individual cycles of the protocol are stitched, aligned, and stacked by the software. The output is a single OME-TIFF image, ready for downstream analysis.

The basic operating principle of the fluidics inside each stainer of the platform is recapitulated in Figure 1c. Each stainer includes a microfluidics-based chip that allows the reduction of incubation times from a typical value of more than 1 hour to less than 5 minutes [22, 25]. The microfluidic chip is placed on the tissue slide loaded onto the staining unit of the instrument. The slide and the chip create a reaction chamber thanks to a seal ensured by a gasket, delimiting the staining area, as well as a mounted piston to clamp the chip onto the slide. Reagents flow through the inlet into the chamber surrounding the sample before exiting through the outlet. The temperature inside the chamber is controlled by a heating/cooling system using a Peltier element placed below the sample slide. The low chamber height (50 µm) and the resulting surface-to-volume ratio enable a rapid reagent exchange to the chamber (less than 1 second) and a rapid temperature control. The chamber temperature can reach 50°C in less than 30 seconds and cool down to room temperature in a period of less than 5 minutes [24, 28].

Once the rotary stage moves a stainer-sample module under the integrated microscope, the setup shown in Figure 1d is formed. COMET™ includes an integrated LED-based widefield microscope that acquires images in three distinct fluorescent channels (DAPI, TRITC, and Cy5) while respective filters prevent crosstalk between channels. The 20x objective enables subcellular resolution of 0.45μm with an image pixel size of 0.23μm. The final mosaic image at full scale is composed of 357 (17×21) tiles, registered and overlayed, creating a single ome.tiff image compatible with various proprietary and open-source software solutions for downstream analysis. At maximum area size, bit depth, and resolution, each layer or channel of the final image requires about 4GB. However, image size can be reduced considerably by choosing different compression options.

### Fully Automated 40-plex seqIF panels on FFPE tissue sections

The seqIF approach on COMET™ was used to develop a 40-plex panel on FFPE human tonsil tissue. The panel consisted of biomarkers inherent to the immune-oncology field: αSMA, BCL6, CD3, CD4, CD8, CD11b, CD11c, CD14, CD15, CD16, CD20, CD21, CD31, CD34, CD38, CD45, CD45RA, CD45RO, CD56, CD68, CD107a, CD138, CD163, panCK, E-Cadherin, FOXP3, Granzyme B, HLA-DR, ICOS, IDO-1, Ki-67, LAG-3, NaKATPase, PD-1, PD-L1, Podoplanin, S100, Tryptase, Vimentin, and VISTA (Table 1). In this panel, markers to identify both the lymphoid (T cells, B cells, and natural killer cells) and myeloid lineage (such as neutrophils, dendritic cells, mass cells, basophils, and macrophages) of the immune system are selected. Complementary to lineage-specific markers, we have selected additional markers to characterize specific functional conditions, such as immunosuppressive status (IDO1, PD-L1, and VISTA), activation of immune cells (PD1, ICOS, and LAG-3), or proliferative status (Ki-67). In addition to the immune profile of the TME, cancer and stromal cells, as well as blood and lymphatic vasculatures (CD31, CD34, Podoplanin) can be detected by specific markers in the panel. The panel also includes markers to study tumors with an epithelial origin (i.e., carcinomas). Tumoral or healthy epithelial cells can be detected by the expression of cytokeratins (CK) or E-cadherin.

**Table 1:**
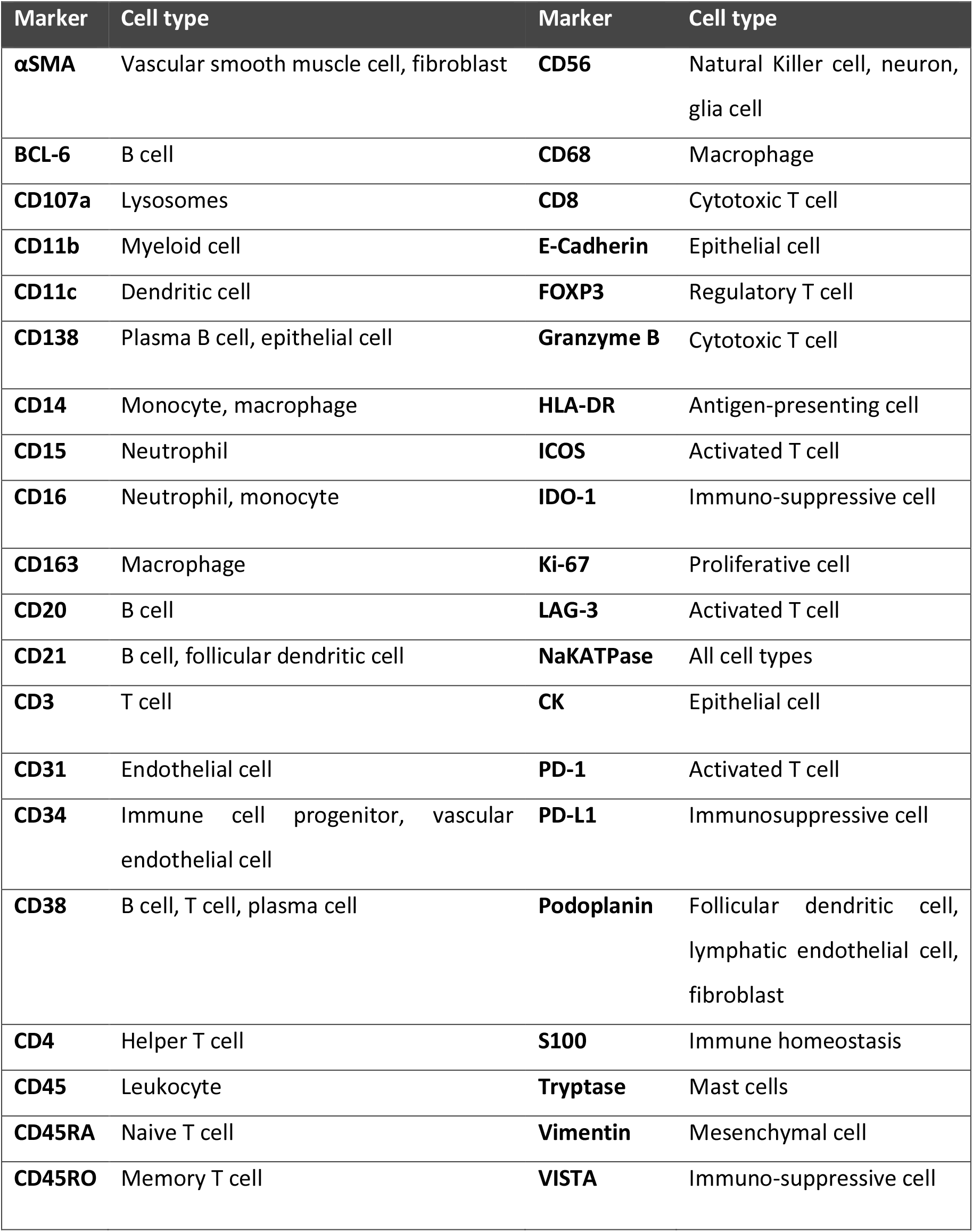
Makers stained for the 40-plex panel and their associated cell phenotypes.

For each marker in the panel, seqIF produced high-quality stainings with satisfactory background to signal ratio and reflecting expression patterns reported elsewhere [29]. Figure 2a shows representative monochrome images from single marker stainings for each marker in the panel and Figure 2b shows the same region of interest with 8 composite overlays, each comprising 5 markers from the panel. These results have been obtained in 20 consecutive cycles of seqIF, by using primary antibody cocktails from 2 species in each cycle. The total assay time was less than 20 hours for the whole run, including the image acquisition step. Performing multiplex immunofluorescence staining with high numbers of markers in one workflow paves the way for an in-depth understanding of the tissue analyzed while drastically reducing the number of samples needed.

**Figure 2:**
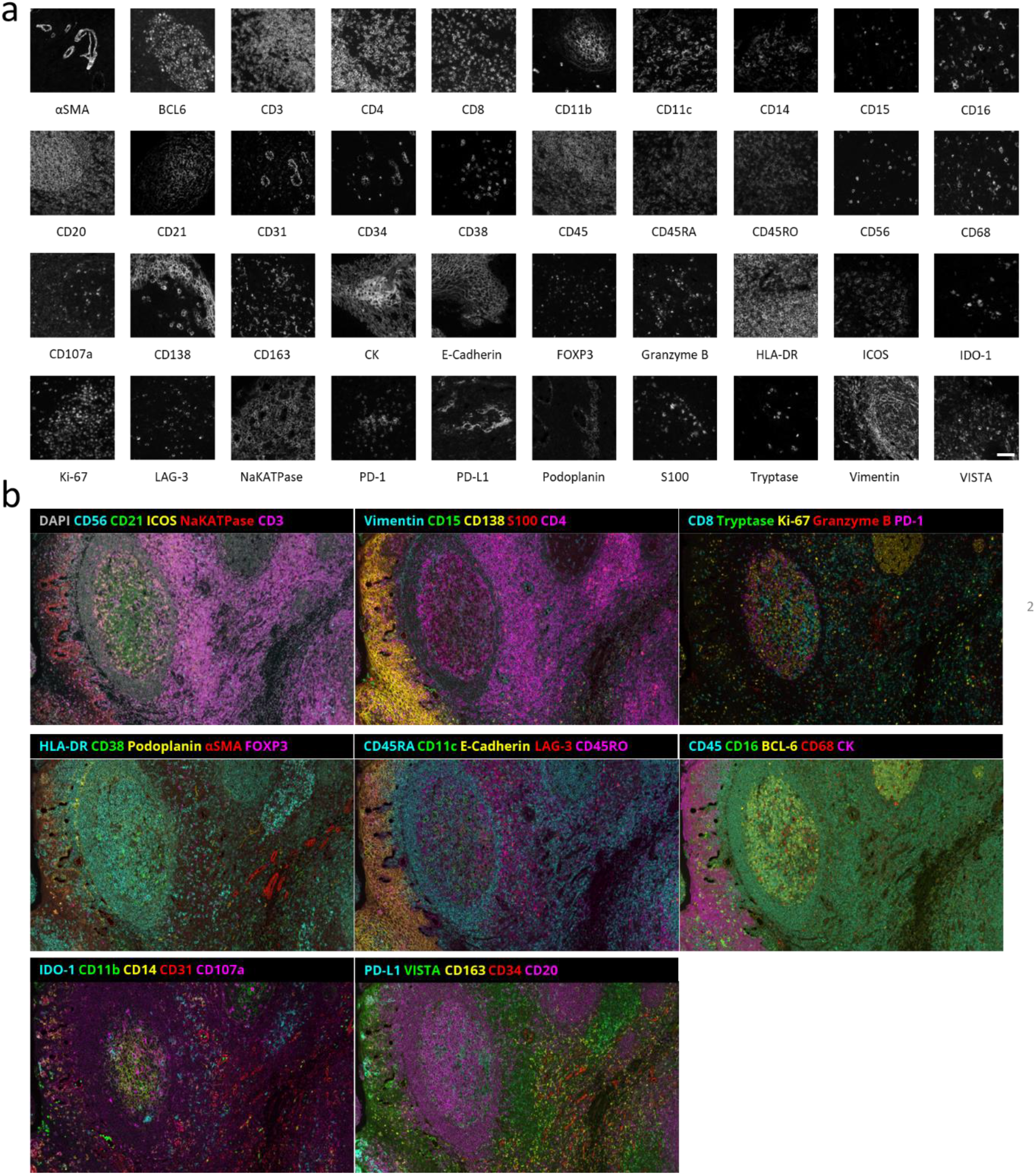
40-marker multiplex staining performed using the seqIF method. **(a)** Snapshots of individual markers from a 40-plex panel staining from human tonsil FFPE samples performed with seqIF on COMET™. Scale bar: 50μm. **(b)** Multiple composite images of a 40-plex staining showing the same region of interest (ROI) of a human tonsil FFPE sample. Autofluorescence subtracted and brightness adjusted for visualization. Scale bar: 100μm.

### Elution Efficiency and Epitope Stability

As seqIF is a technique based on stripping primary and secondary antibody complexes off the tissue in between each staining cycle, elution represents a critical step in the protocol. Antibody elution must be efficient in removing antibodies to ensure staining specificity and repeatability for subsequent cycle of staining, but at the same time gentle enough to preserve the tissue integrity and target epitopes over numerous cycles, enabling a reliable detection for multiplex staining. Figure 3a shows elution efficiency and epitope stability analysis results obtained with a dedicated protocol that was run on COMET™ for CD20 staining on human FFPE tonsil tissue. Using the negative controls, where only secondary antibodies are dispensed before and after elution, as well as the staining image, it is possible to calculate the elution efficiency of the marker (Methods and Figure S5). For example, for the CD20 staining depicted in Figure 3a, the elution efficiency was determined as 99.4%. To ensure high elution efficiency, several different tissue types were examined. The elution efficiency studies were run on human FFPE tonsil, human FFPE colorectal cancer, and mouse frozen section samples (1 sample of each type), with 3 markers each, where quantitative analysis via comparison of signal intensity between the stainings and negative controls showed that elution efficiency was greater than 95% in each case (Figure 3b, see the calculation method in supplementary information). Overall, these results highlight robust antibody removal during the elution step, with no tissue degradation observed.

**Figure 3:**
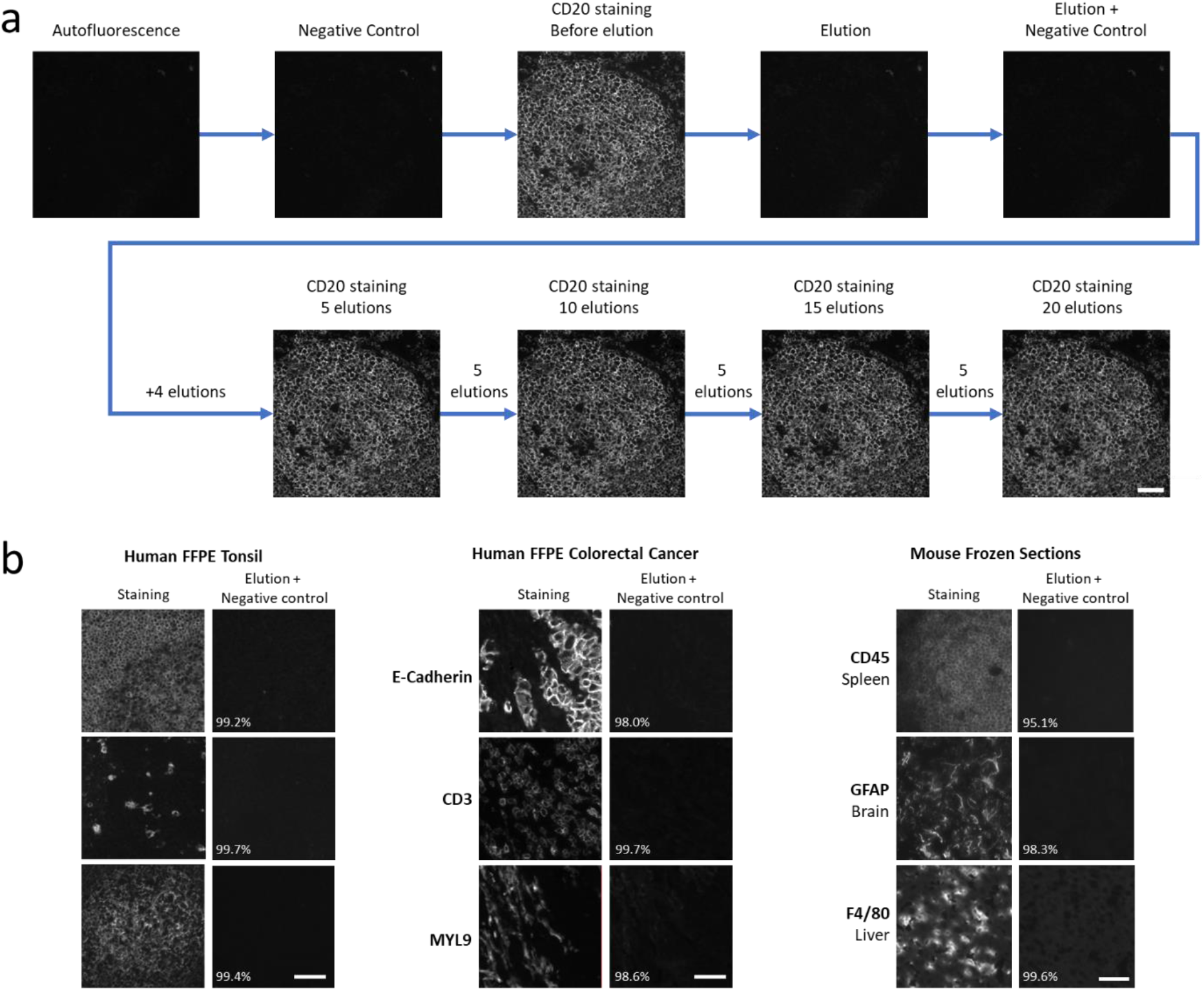
Elution efficiency assessment of seqIF. **(a)** Elution efficiency and epitope stability of CD20 staining on a human tonsil FFPE sample across 20 elution cycles. Scale bar: 50μm. **(b)** Example of elution followed by negative controls on different tissue samples, with elution efficiency indicated as a percentage. The first panel shows the elution efficiency of the markers CD45, CD68, and CD19 on a human FFPE tonsil sample. The second panel shows the elution efficiency of the markers E-Cadherin, CD3, and MYL9 on a human FFPE colorectal cancer sample. The third panel displays the elution efficiency of CD45, GFAP, and F4/80 on different mouse frozen section tissues (spleen, brain, and liver). Scale bars: 50μm.

An important parameter in building mIF panels is the optimal placement of each marker in terms of its order in the protocol. To assess the position of a biomarker in a seqIF protocol, it is essential to evaluate the detection of the target tissue epitope over multiple protocol cycles. This approach is particularly valuable in bigger panels (e.g., greater than 20-plex) because of the increased number of steps that tissue undergoes during the run. To this end, we used the second part of the protocol depicted in Figure 3a, consisting of intercalating multiple cycles of elution between staining steps, to compare the initial staining to subsequent stainings for up to 20 elution cycles. The guided assay optimization approach on COMET allows two different modes to assess biomarker stability: (i) A high number of cycles (e.g., 20) can be executed to evaluate staining quality after each elution or (ii) Elution cycles can be merged, and images can be acquired after 5 consecutive rounds of elution. The latter allows to decrease the total protocol time, reagent consumption, and dataset file size, while the former enables an increased accuracy of epitope stability determination (Figure S3a). At constant intervals of 5 cycles, we then evaluated the marker epitope stability by comparing the signal to background ratio (SBR) between the image obtained in the first cycle and the subsequent images registered after 1, 5, 10, 15 and 20 elution cycles (Figure 4). For all the 4 markers analyzed, we have observed a negligible drop of SBR, demonstrating high epitope stability, with panCK showing the highest degree of variations (5.6% drop after the 5^th^ elution). However, the SBR values all remain constant after subsequent elution cycles, demonstrating the performance of the seqIF method for the preservation of epitope stability. Using this approach to characterize the behavior of each biomarker, we can also deduce that if a marker displays a significant drop of SBR after several elution cycles, it should be placed in the early cycles of the multiplex protocol, while a marker with an excellent epitope stability can be placed in any cycle of the protocol, meaning at any position in the panel. In overall, the data presented here highlights the robust epitope preservation capability of seqIF protocols that enables flexibility while designing multiplex panels.

**Figure 4:**
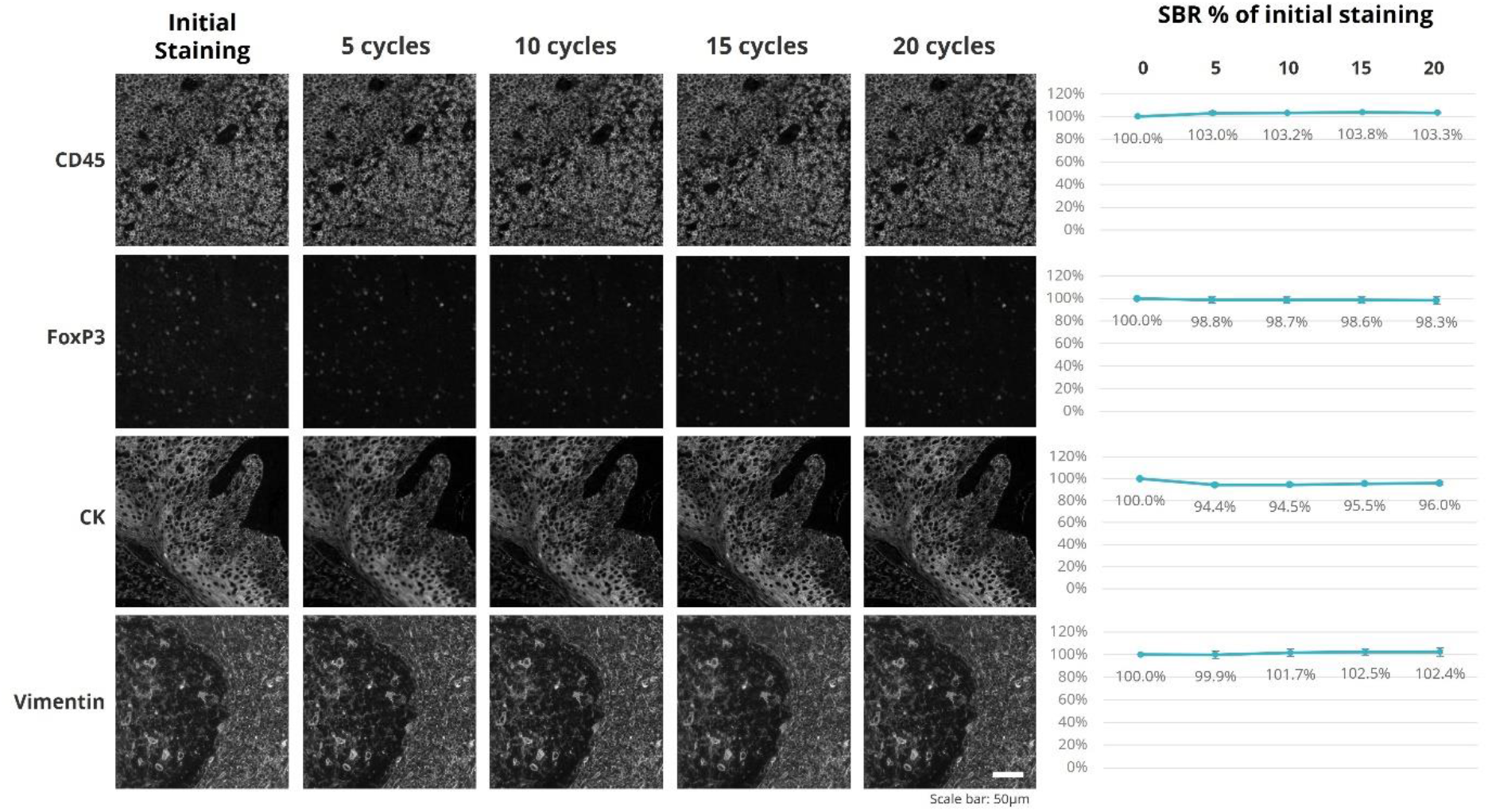
Staining intensity evaluation over 20 iterations of seqIF protocol. The epitope stability of four markers targeting different cell types and cellular localization (CD45, FoxP3, panCK, and Vimentin) quantified after 5 cycles, 10 cycles, 15 cycles and 20 cycles of elution compared to the initial staining. All staining and elution steps were performed on the same tissue sample (human tonsil FFPE). The signal-to-background ratios (SBR) as a % of the initial staining are displayed on the right. Scale bar: 50μm. n=1.

### Repeatability and Reproducibility

To assess the robustness of seqIF protocols on COMET™, we have run intra-site repeatability and reproducibility studies for a panel of 4 markers on consecutive slides of human FFPE tonsil tissue, staining different cell types and compartments (FoxP3 presents in nuclei of regulatory T cells, CD68 expressed in cytoplasm of macrophages, CD3 present on T cells membranes, and CD20 marking B cells membranes). The following aspects were considered in the repeatability and reproducibility studies: (i) To evaluate if the protocol performance is consistent among consecutive runs, consecutive run repeatability studies were performed (same device, same day, and same stainer) (Figure 5a), (ii) To investigate if the staining performance is impacted by the stainer unit used on the instrument, repeatability on different stainer units was evaluated (same device and same day) (Figure 5b), (iii) Consecutive day reproducibility tests were run (same device) to determine if reproducible results can be obtained over several days on the same instrument (Figure 5c), and finally (iv) Different device reproducibility tests were carried out (run on the same day) to verify if different instruments have comparable staining and imaging performance (Figure 5d). The results were quantitatively assessed by calculating the coefficient of variation (CV) of the SBR for each marker (see Methods). The CV was determined to be minimal (<10%) for the majority of the conditions tested, with acceptable performance across the board, where FoxP3 showed considerable CV for the inter-stainer and inter-device experiments (Figure 5). These results show that seqIF combined with the fully automated workflow of COMET™ ensures robustness and reproducibility of obtained results with very low variability.

**Figure 5:**
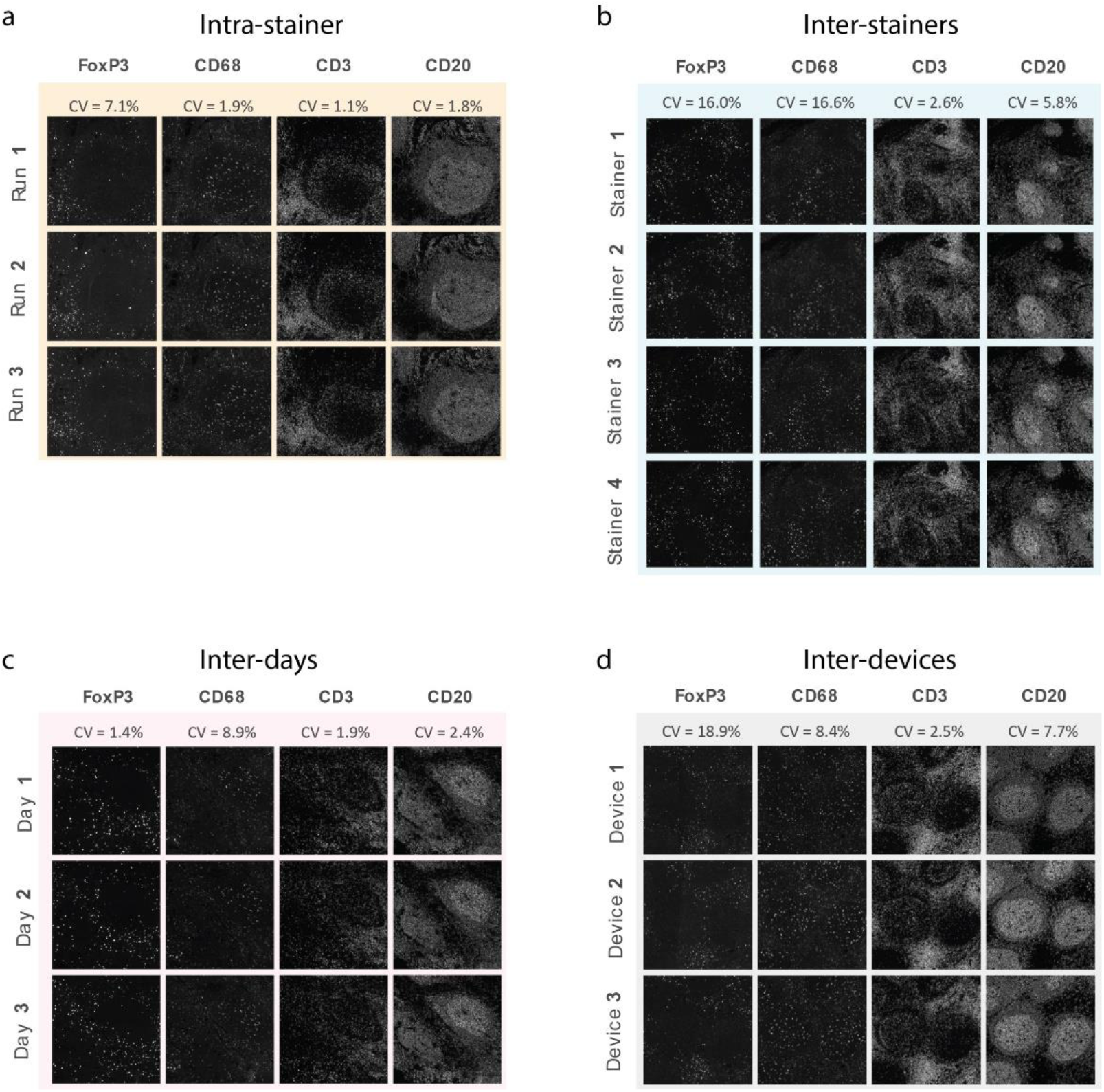
seqIF repeatability and reproducibility studies run for 4 markers on human FFPE tonsil tissue. **(a)** Intra-stainer repeatability was assessed by 3 subsequent runs performed on the same staining unit during the same day and resulted in CVs below 8%. **(b)** Repeatability between independent staining units was assessed by executing the same protocol all stainers of the same COMET™ device and resulted with CVs below 17%. **(c)** Reproducibility of the assay was assessed by the same protocol being executed on the same staining unit of the same device on independent days and resulted in CVs below 10%. **(d)** Reproducibility of the assay between different devices was assessed by execution of the same protocol on different platforms on the same day and resulted in CVs below 20%. CV stands for coefficient of variation of SBR between different images in the dataset. n=1.

### Signal Uniformity and Dynamic Range

In most cases, images obtained through seqIF protocols on COMET™ require further downstream analysis. Signal uniformity is an important parameter to ensure that images that are being analyzed can give consistent and reliable results that reflect biomarker expression patterns. Another important parameter is the dynamic range of the fluorescent signal, which allows the interpretation of the output for a wide spectrum of protein marker expression values. Figure 6 shows examples of signal uniformity and dynamic range assessment on seqIF data sets. The quantification of signal variation for CD20 staining on an FFPE human tonsil tissue shows the variation below 6% for both axes over the whole image (Figure 6a). The wide dynamic range for FoxP3 staining on the same sample shows that low, mid, and high expressing cells can be well discriminated, suggesting that expression heterogeneity is preserved, and the linear relationship between the signal and the antigen amount is preserved (Figure 6b). Another important process to ensure that obtained images can be properly analyzed is the compensation of the tiling effect that occurs after the assembly of final mosaic images from single fields of view (FoV). The tiling effect is linked to light variation and distortion in the optical path, as shown in Figure 6c. To improve the image homogeneity, automatic flat-field correction is applied to every imaged FoV using a flat-field reference image. Our data confirms that seqIF images are of sufficient quality in terms of dynamic range and uniformity to enable in-depth biomarker analysis, including expression level, within the analyzed tissue.

**Figure 6:**
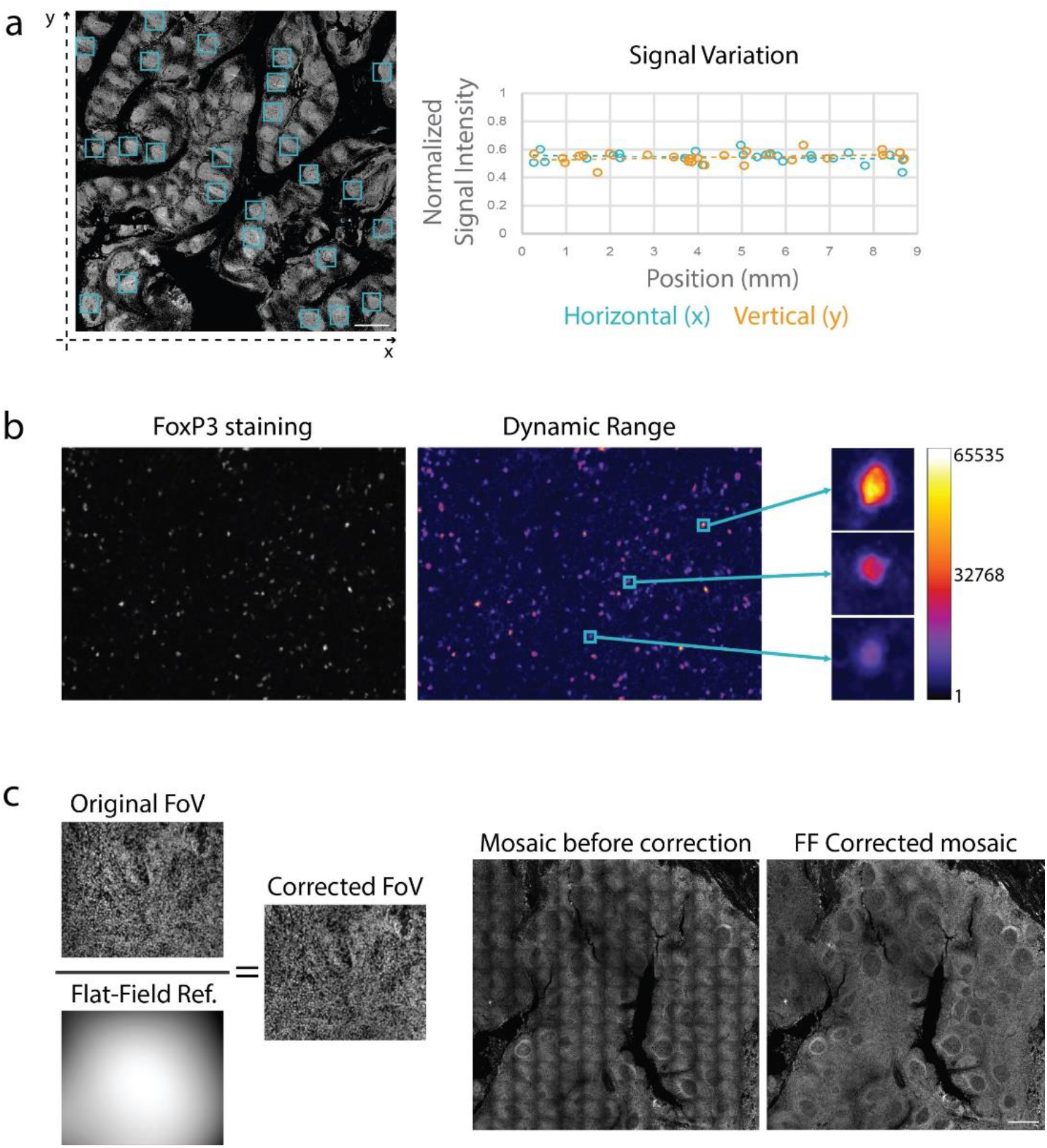
Demonstration of aspects that ensure image quality with seqIF and COMET. **(a)** Quantification of the horizontal and vertical signal variation of a staining against CD20 on human tonsil FFPE performed and imaged on COMET™. The normalized signal intensities of 25 regions of interest (left, blue squares) were plotted according to their horizontal (right graph, blue) or vertical position (right graph, orange). Overall signal variation for both axes is below 6%. Both X and Y values are represented in millimeters. Scale bar: 1 mm. **(b)** Example of the dynamic range of a FoxP3 staining on COMET™ (left) and pseudo coloring with a Fire look up table (right) highlighting low, mid, and high expressing cells. The scale represents grey values of the 16-bit images (1 to 65535). **(c)** Automated flat-field correction provides uniform seqIF image. The result is a seamless mosaic of the whole image without aberrations (right) compared to an image without correction and visible aberrations (left).

### Single Cell Analysis on seqIF Images

Data sets obtained with seqIF are suitable for single cell analysis (Figure 7). We present here results from a 10-plex panel run on an FFPE human tonsil tissue sample. The multi-stack OME-TIFF output was visualized, and brightness settings were adjusted for each marker using the Lunaphore Viewer software, a visual tool fully integrated in the staining platform for the qualitative assessment of staining results. The upper panel in Figure 7a shows a caption on the germinal center border, with a 3-marker subset of the original 10-plex panel shown (CD20: cyan, CD3: magenta, CK: yellow). The bottom panels display three zoomed-in regions in which the combination of markers allows the identification of multiple cell subtypes: Epithelial cells (CK^+^, yellow), T cells (CD3^+^, magenta) and B cells (CD20^+^, cyan) on the bottom left, macrophage cells (CD11c^+^CD4^+^) and proliferating T helper cells (CD4^+^Ki-67^+^CD11c^-^) on the bottom center, and immature T cells (CD4^+^CD8^+^), T helper cells (CD4^+^CD8^-^), T killer cells (CD8^+^CD4^-^) and T regulatory cells (CD4^+^FoxP3^+^CD8^-^) on the bottom right. An important aspect linked to the analysis is the autofluorescence subtraction, which can serve as an aid for the visualization of certain markers (Figure 7b). This holds relevance for tissues with high intrinsic fluorescence where specific staining signal intensity may be at similar levels as the tissue autofluorescence, making the distinction of the specific signal cumbersome. seqIF and COMET™ provide standardized options to use this feature where needed. Figure 7b shows the endogenous tissue autofluorescence (magenta) and CD68 staining (green) of a human lung adenocarcinoma FFPE sample. Without subtraction of the autofluorescence, the true CD68 signal is hard to distinguish from the bright tissue-intrinsic autofluorescence, while the background subtraction allows the better appreciation of the CD68 positive cells.

**Figure 7:**
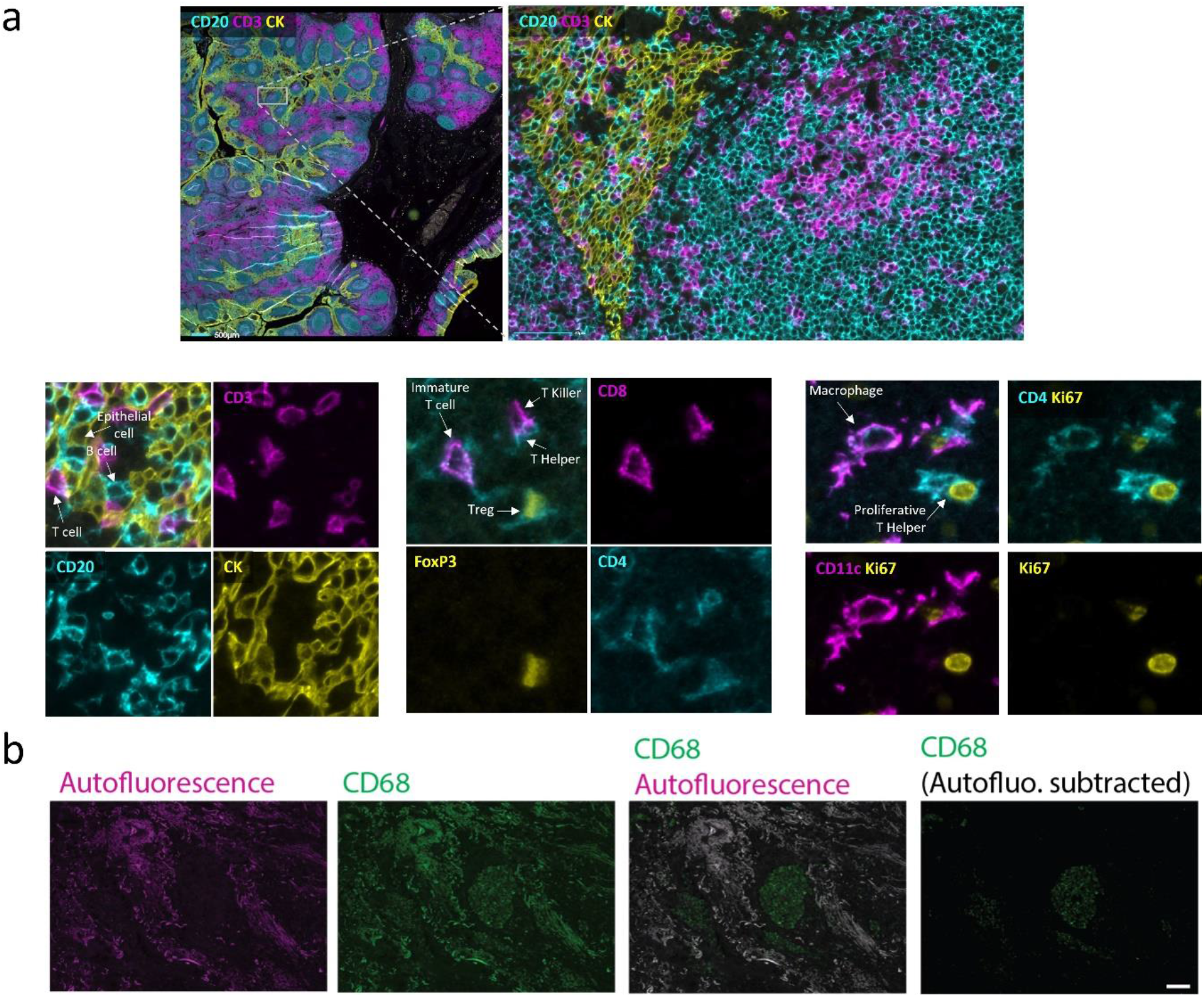
seqIF images on COMET provide subcellular resolution for single cell analysis. **(a)** Upper panel: Border of a human tonsil, germinal center FFPE sample, with a subset of a 10-plex panel staining (CD20: cyan, CD3: magenta, CK: yellow). Scale bars: 1mm (left) and 100µm (right). Lower panels: Three zoomed-in regions where multiple cell subtypes are identified, including epithelial cells, B cells, T cells (bottom left). Macrophage cells and proliferating T helper cells (bottom center). Immature T cells, T helper cells, T killer cells and T regulatory cells (bottom right). Scale bar: 10µm. **(b)** Endogenous tissue autofluorescence (magenta) and CD68 staining (green) of a human lung adenocarcinoma FFPE sample. Scale bar: 100µm.

## Conclusion

The landscape of emerging technologies in the field of spatial biology is evolving at a fast pace [10] with a constantly increasing number of high-plex platforms intended to reveal in-situ biomarker expression in a variety of clinical contexts [30]. Many of these platforms exploit different detection approaches to spatially identify multiple biomarkers in a single tissue section, ranging from using iterative steps of staining and imaging followed by chemical or enzymatic release of macromolecular complexes, to laser or ion beams coupled to mass spectrometers to analyze antibody-metal atom conjugates, spatially indexed beads, and oligo-barcoded technologies to look at different molecular entities [31–33]. While this landscape variety provides researchers with a valuable toolkit to visualize and dissect intricate details of different biological contexts and determine the dynamics of cell interactions at a single-cell level, there is still an increasing need for highly resolved spatial multiplex biomarker detection on tissue samples.

We developed a fully automated sequential immunofluorescence method that is highly suitable to address this need. The technique, based on COMET™ technology, enables lower turnaround times compared to alternatives for multiplexed sample staining thanks to improved reaction dynamics in the microfluidic chamber. Further, in-situ imaging coupled with gentle elution-based signal removal results in well preservation of sample morphology over numerous cycles in seqIF assays, thus higher plex levels. The in-situ imaging approach also allows the process to be streamlined, with walk-away automation resulting in a “sample-in, image-out” workflow. Overall, the developed seqIF method allows a 40-plex assay to be run and results obtained within less than 24 hours. It should also be noted that the method would allow the plex level to be increased to much higher numbers by increasing the number of seqIF iterations or using additional channels to detect more biomarkers per cycle.

SeqIF is a versatile tool for use in different biological domains, including both basic research and translational science. The ability to characterize cell types and the further classification of sub-phenotyping also makes it suitable for specific areas like immune oncology. For example, currently one of the most important use areas of seqIF is the investigation of the TME composition and response to stimuli, where the characterization of cell types and the identification of subtypes play a crucial role in understanding the underlying biological processes. This use is expanded further through the analysis of disease or tissue specific markers such as transcription factors. We foresee that with the increased adoption of spatial proteomics in the research community, seqIF will become a part of established -omics practices, and its application areas will continue to naturally expand, eventually becoming a staple stone of translational research, with a strong outlook towards the clinic.

## Methods

### Ethical Approval

This study has been approved by Commission cantonale (VD) d’éthique de la recherche sur l’être humain (CER-VD), project ID: 2022-01489.

### Materials

Lunaphore MK03 imaging chips have been used with the automated staining and imaging platform COMET™ for all immunostaining experiments. Generic reagents used during the staining and washing steps are as follows: De-ionized water, Alcohol 70%, Multistaining Staining Buffer (Lunaphore BU06), Elution Buffer (Lunaphore BU07-L), Quenching Buffer (Lunaphore BU08-L), Imaging Buffer (Lunaphore BU09), Blocking Buffer (Lunaphore BU10).

Primary antibodies, along with their corresponding protocol conditions, including concentration and incubation time, are listed in Supplementary Table 1.

The used secondary antibodies and counterstaining solution are as follows: Goat anti-Mouse IgG (H+L) Highly Cross-Adsorbed Secondary Antibody, Alexa Fluor™ Plus 555, Goat anti-Mouse IgG (H+L) Highly Cross-Adsorbed Secondary Antibody, Alexa Fluor™ Plus 647, Goat anti-Rabbit IgG (H+L) Highly Cross-Adsorbed Secondary Antibody, Alexa Fluor™ Plus 555, Goat anti-Rabbit IgG (H+L) Highly Cross-Adsorbed Secondary Antibody, Alexa Fluor™ Plus 647, Goat anti-Chicken IgY (H+L) Cross-Adsorbed Secondary Antibody, Alexa Fluor™ Plus 555, Goat anti-Rat IgG (H+L) Highly Cross-Adsorbed Secondary Antibody, Alexa Fluor™ Plus 647 (Thermo Fisher Scientific), DAPI Solution (Thermo Fisher Scientific).

### Tissue Sections

Whole section cuts of formalin-fixed, paraffin-embedded (FFPE) human tonsil, human lung adenocarcinoma, human colorectal cancer and human breast carcinoma tissues were used in the experiments to generate the data shown in this study (obtained from Tissue Biobank Bern). Additionally, different mouse frozen section tissue samples (spleen, brain, and liver) were used in elution efficiency assessment experiments (Amsbio).

### Human FFPE Tissue Pre-processing

Prior to sequential immunofluorescence, FFPE slides were pre-processed through dewaxing and antigen retrieval processes. The dewaxing and antigen retrieval were performed on Epredia Lab Vision™ PT Module instrument (Epredia), using Dewax and HIER (Heat Induced Epitope Retrieval) Buffers (Epredia) at pH 9. The incubation was performed for 60 minutes at a temperature of 102°C. The processed slides were then transferred to a Multistaining Buffer (Lunaphore, BU06) bath until use.

### Mouse Frozen Section Pre-processing

Frozen section (FS) slides were air-dried for 20 min at room temperature prior the incubation in 4% formaldehyde (Sigma Aldrich) on an orbital shaker for 40 min. Subsequently, slides were washed for 5 min in Multistaining Buffer (BU06, Lunaphore Technologies) and incubated in 0.2% Triton in Staining Buffer (BU01, Lunaphore Technologies) for 20 minutes on an orbital shaker. Subsequently, slides were stored in Multistaining Buffer (Lunaphore, BU06) until use.

### Signal Intensity, Coefficient of Variation (CV) and Elution Efficiency Calculations

For the repeatability and reproducibility analysis, the mean signal intensity was calculated from 6 different regions of interest of 1 mm^2^ using Otsu thresholding for pixel-based analysis [34]. The coefficient of variation (CV) of the mean signal intensity was used in experiments where the variation between images was measured.

The elution efficiency calculations for the characterization of protocols developed in this study were based on the following equation:

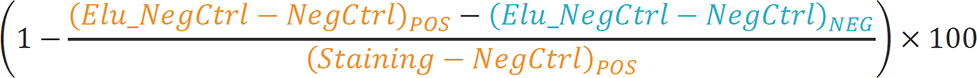

Here, we evaluated the elution efficiency as the efficiency of staining removal in the positively stained areas upon elution, expressed as the complement of the percentage of the difference in intensity between the negative control signal after elution and negative control signal before staining in the positively stained area, with respect to the intensity of the staining in the same area (Figure S5). In order to measure the effective elution efficiency of the staining, independently from general background changes due to the elution cycle, we subtracted the difference between negative control signal after elution and negative control signal before staining in the negative, unstained area (Figure S5).

## Data availability

The datasets used and/or analyzed during the current study available from the corresponding author on reasonable request.

## Author Contributions

F.R., J.K., A.K., M.G.P., P.B., E.P. and S.B. conceived, executed, and analyzed experiments. D.E., B.P., M.A., E.D., S.S., M.R., G.CE., P.J. and D.D. contributed to platform design, development, and testing. L.B., J.F., G.CO. and L.A. contributed to platform software development. M.C. and D.D. supervised the manuscript process. D.E. wrote the manuscript. F.R. composed the figures. F.R., J.K., M.C., B.P. and D.D. reviewed the manuscript.

## Notes

### Competing Interest Statement

All authors of the manuscript are current or former employees of Lunaphore Technologies SA, which is commercializing an automated version of the seqIF method.

