## Supplementary Information for "Fully Automated Sequential Immunofluorescence (seqIF) for Hyperplex Spatial Proteomics"

#### *Qualitative Assessment of Repeatability and Reproducibility*

To assess the repeatability and reproducibility of the developed method further, a 10-plex panel (CD68, FOXP3, CD8, PD-1, Ki-67, PD-L1, CK, CD4, CD20, CD3) on human tonsil FFPE tissue was run on 4 stainers of a COMET™ instrument in a single run (Figure S1a), as well as three times in a row on the same stainer (Figure S1b). Qualitative assessment of the results showed no visible variability on the obtained results.

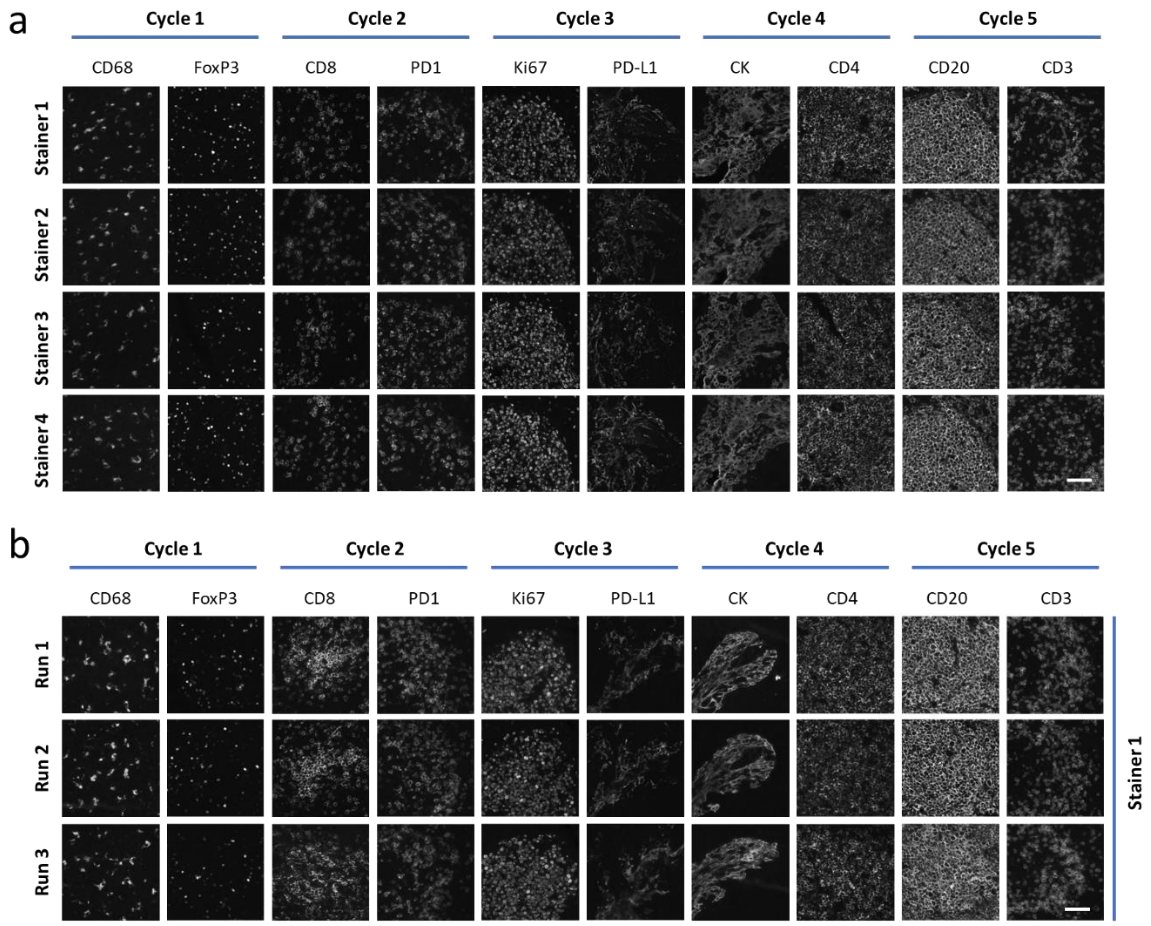


**Figure S1:** seqIF assays on COMET™ show high reproducibility between different stainers and independent runs.

**(a)** seqIF inter-stainer reproducibility study results on COMET™. A 10-plex panel protocol was run on the 4 stainer units of the same device and on the same day. Each image displays a representative ROI of a single marker. Scale bar: 50 µm.

**(b)** seqIF intra-stainer repeatability study results on COMET™, where consecutive runs of the same 10-plex protocol on multiple slides were executed on the same stainer unit of the same device. Scale bar: 50 µm.

#### *User Workflow Overview for SeqIF*

Figure S2 shows a broader view of the used seqIF workflow. It starts with sample pre-processing, which is performed in a PT Module. In parallel, device initialization, reagent preparation and loading of the reagents, protocols, and slide samples onto COMET™ is done. Once the slides are loaded in the instrument, the fully automated seqIF protocol starts. When needed, the method also allows the inclusion of pauses in the protocol to adjust imaging parameters in between cycles. This is a step that is typically needed during marker optimization or panel development, but to a lower extent in higher throughput experiments such as batch stainings since all parameters including imaging would be optimized beforehand. Considering that the developed method has a typical duration of 30 minutes per cycle, and 2 markers per cycle can be stained and imaged by using mixes of antibodies from different species, the process time for a typical 40-plex run is in the order of less than one day.

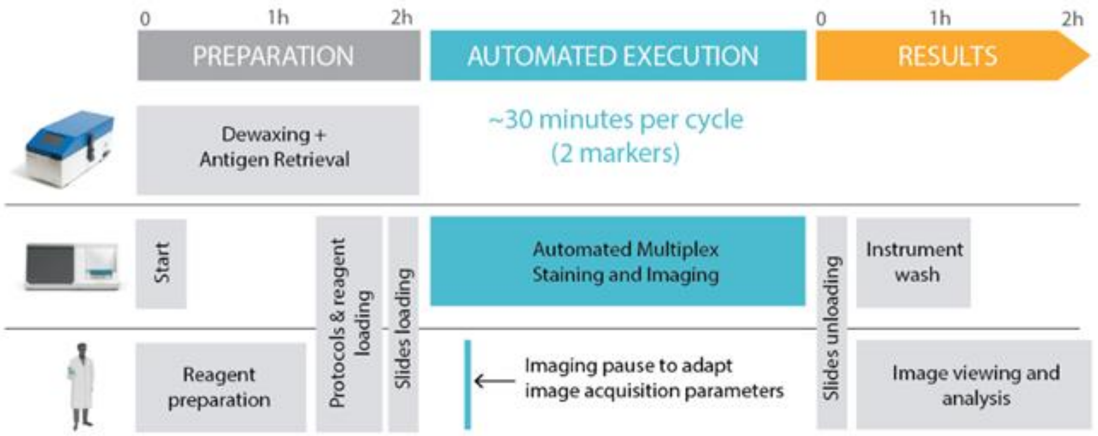


**Figure S2:** Overview of seqIF workflow, including timeframes for reagent and sample preparation, seqIF execution by automated platform and analysis of results.

### ***Biomarker and Panel Optimization in seqIF***

We have developed multiple seqIF marker and panel optimization strategies and characterization techniques to obtain results in the fastest time frame possible, and with the least amount of sample use. Figure S3a shows a representation of the developed approaches implemented in the form of different guided protocol templates available on COMET™. "Characterization Part 1" is a protocol that is used to assess the staining quality and elution efficiency of a new antibody prior to any optimization. It consists of a single staining cycle with imaging performed before and after the initial staining as well as after the elution step. An optional negative control step can be integrated after the elution. "Characterization Part 2" is designed to test different parameters on multiple cycles to determine optimal staining conditions. Images are acquired both after the staining and the elution steps. The user can set up to 20 cycles with different conditions. "Characterization Part 3" assesses the elution efficiency of a staining and its associated epitope stability over up to 20 cycles. SeqIF is the main protocol to run previously optimized multiplex staining protocols, for up to 20 cycles.

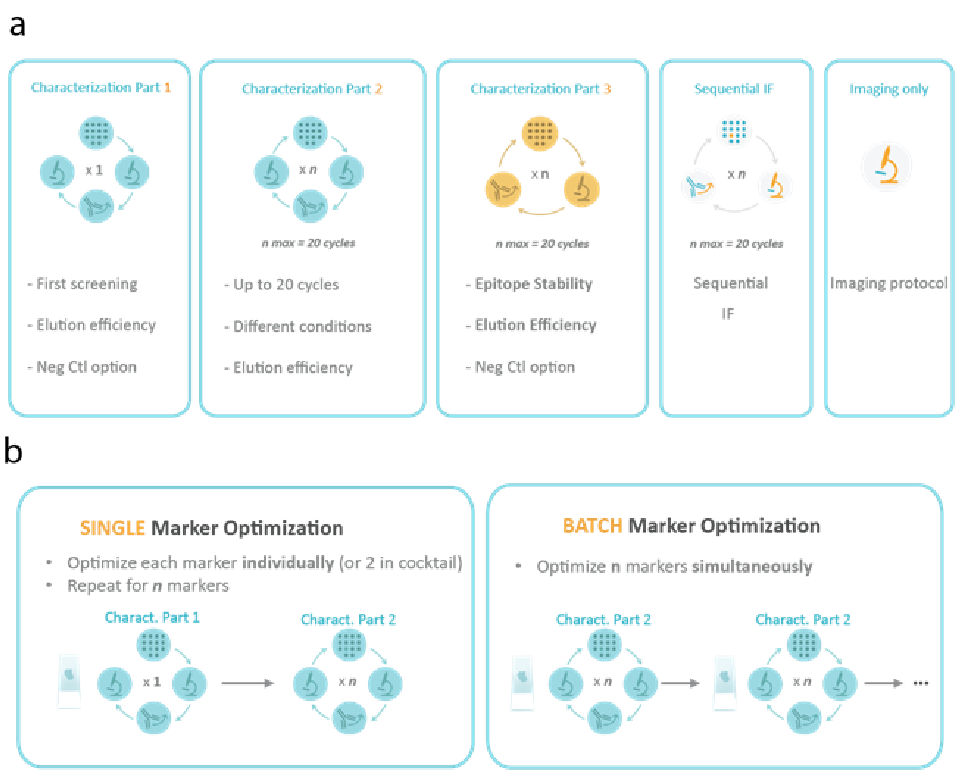


**Figure S3**: Guided optimization approaches on COMET™ for seqIF protocol development.

**(a)** Assisted optimization protocols developed for seqIF.

**(b)** Single and batch marker optimization modes used in panel development and optimization on COMET™. Neg Ctl: Negative Control - Tissue incubation with only secondary antibodies or with isotype controls of primary and secondary antibodies.

The two main optimization approaches for seqIF, single-marker and batch optimization modes, are shown in Figure S3b. During single marker optimization, up to 2 primary antibodies can be optimized for different conditions at each cycle using an initial Characterization Part 1 protocol, followed by Characterization Part 2 to test more conditions on the same tissue slide. If more than 2 markers require optimization, batch marker optimization becomes a useful approach. When batch optimization is preferred, multiple markers and conditions are tested in the same experiment using Characterization Part 2 protocols on consecutive tissue samples and optimize the desired number of markers simultaneously.

Figure S4 shows an example from the batch optimization approach where several markers are run in a relatively small panel, and parameters are adapted for each new run on a human lung adenocarcinoma FFPE sample on COMET™. The IHC references were provided by the tissue biobank. The 10-plex panel displayed was run as a 5-cycle protocol on a single tissue sample slide. Primary antibodies raised in different host species were mixed and detected by their corresponding secondary antibodies at each cycle. Using serial slides for each marker, a total of 4 runs were necessary for this panel optimization. Between each run, antibody concentration adjustment, primary antibody clone change, or the use of a blocking solution (blue arrows) were performed to reach the optimal final conditions in run #4.

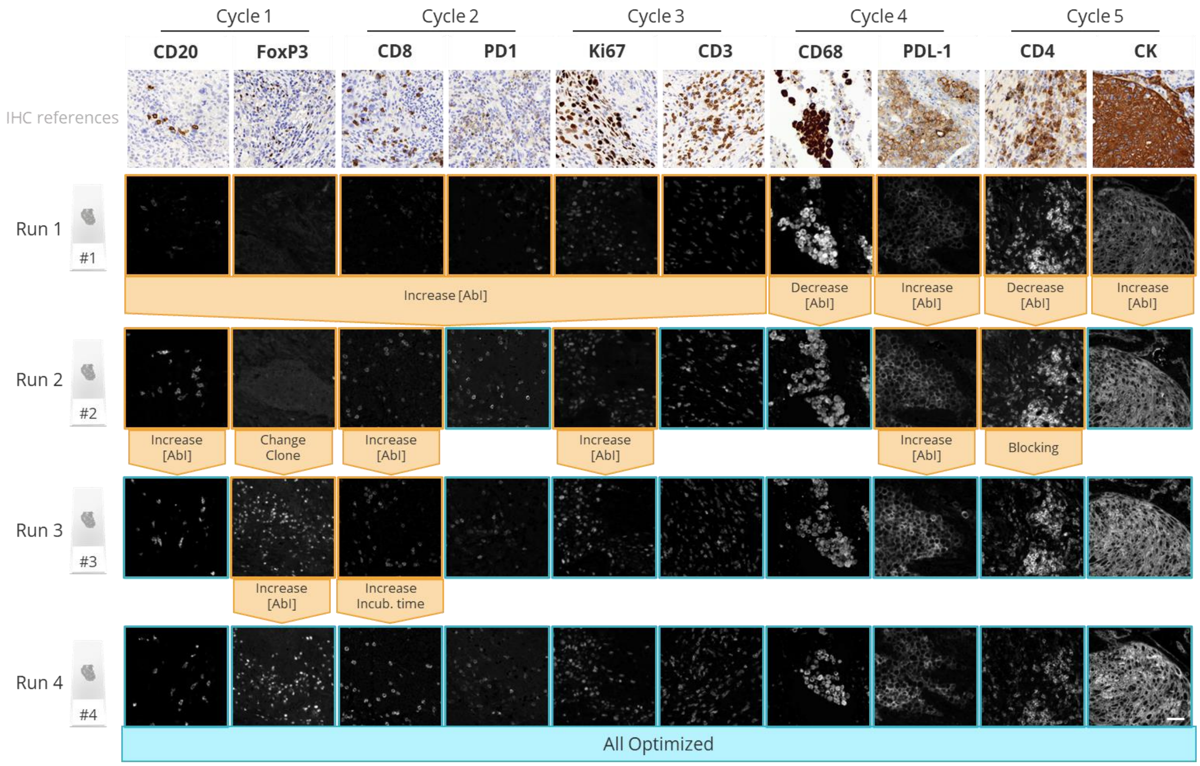


**Figure S4:** Batch marker optimization example. Optimization of a protocol performed on a human lung adenocarcinoma FFPE sample on COMET™, showing optimized results reached after 4 runs for a 10-plex panel.

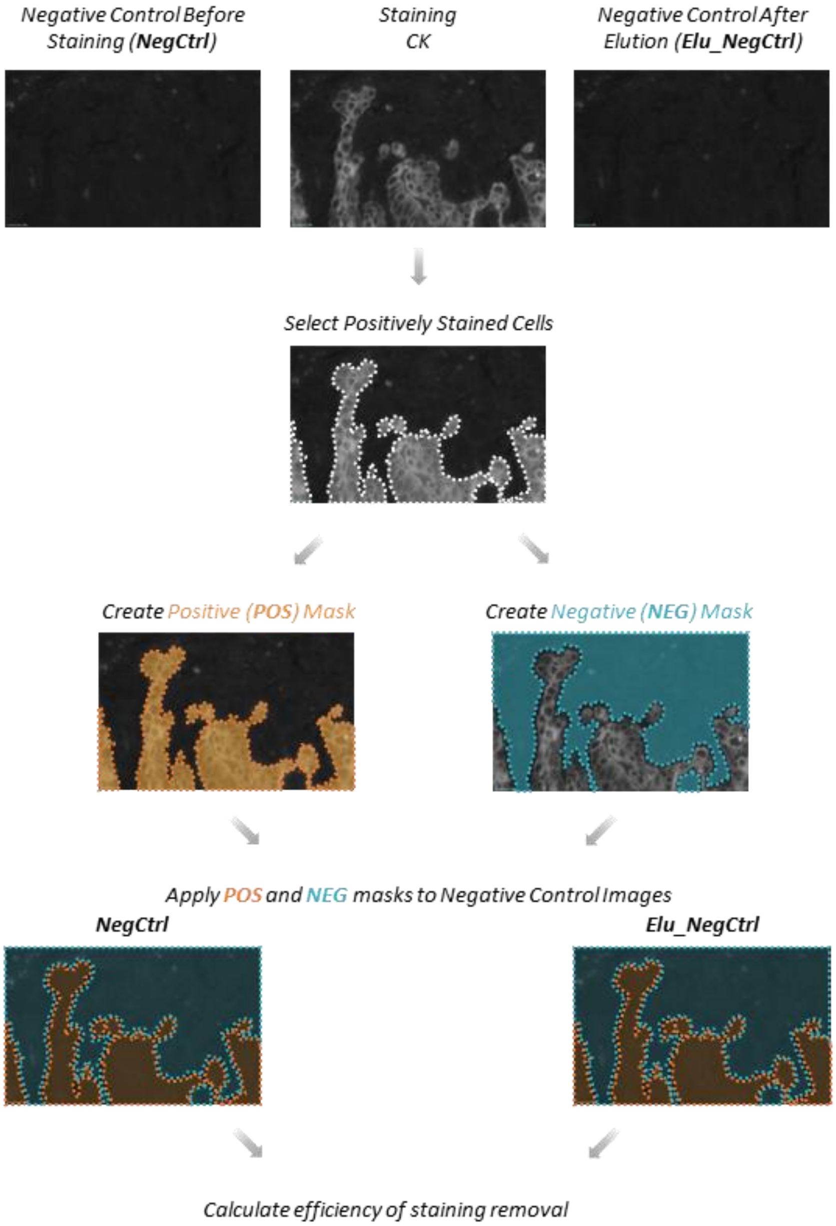


**Figure S5:** Demonstration of elution efficiency calculation workflow, where negative control images before staining and after elution are used in combination with the staining image to obtain the efficiency of staining removal.

NegCtrl: Negative Control - Tissue incubation with only secondary antibodies or with isotype controls of primary and secondary antibodies

Elu_NegCtrl: Negative Control after elution - Tissue incubation after elution of staining with secondary antibodies only.

**Table S1**: Primary antibody list and conditions for seqIF protocol.

| Antigen | Antibody clone | Antibody Species | Supplier | Dilution range (1/) | Incubation time (minutes) |
|---|---|---|---|---|---|
| αSMA | 1A4 | Mouse | Cell Marque | 200-400 | 4 |
| BCL6 | EP278 | Rabbit | Cell Marque | 50 | 4 |
| CD107a | H4A3 | Mouse | Biolegend | 100 | 4 |
| CD11b | EPR1344 | Rabbit | Abcam | 800 | 4 |
| CD11c | EP157 | Rabbit | Cell Marque | 100 | 4 |
| CD11c | EP1347Y | Rabbit | Abcam | 1600 | 4 |
| CD138 | MI15 | Mouse | Biolegend | 100 | 4 |
| CD14 | EPR3653 | Rabbit | Abcam | 200-1000 | 4 |
| CD15 | CARB-3 | Mouse | Dako | 500 | 4 |
| CD16 | SP175 | Rabbit | Cell Marque | 100-400 | 4 |
| CD163 | MRQ-26 | Mouse | Cell Marque | 50 | 4 |
| CD19 | EPR5906 | Rabbit | Abcam | 200-400 | 4 |
| CD20 | L26 | Mouse | Cell Marque | 100-200 | 4-8 |
| CD20 | SP32 | Rabbit | Thermo Scientific | 100 | 4-8 |
| CD21 | EP3093 | Rabbit | Cell Marque | 400-1000 | 4 |
| CD3 | MRQ39 | Rabbit | Cell Marque | 200-1000 | 4-8 |
| CD31 | EPR3095 | Rabbit | Abcam | 400-800 | 4 |
| CD34 | EP88 | Rabbit | CellMarque | 400 | 4 |
| CD38 | SP149 | Rabbit | Cell Marque | 400-800 | 4 |
| CD4 | EPR6855 | Rabbit | Abcam | 100-400 | 4-8 |
| CD45 | PD7/26 + 2B11 | Mouse | Dako | 50 | 4 |
| CD45 | 30-F11 | Rat | Cell Signaling | 1000 | 2 |
| CD45RA | HI100 | Mouse | BioLegend | 1000-1600 | 4 |
| CD45RO | UCHL1 | Mouse | Dako | 200 | 4 |
| CD56 | MRQ42 | Rabbit | Cell Marque | 100-600 | 4 |
| CD68 | KP1 | Mouse | Thermo Scientific | 50-100 | 4 |
| CD8 | 4B11 | Mouse | BioRad | 50-100 | 4-8 |
| CK | AE1/AE3 | Mouse | Dako | 50-150 | 4 |
| E-Cadherin | 36/E | Mouse | BD Bioscience | 800-1000 | 4-8 |
| F4/80 | BM8.1 | Rat | Cell Signaling | 400 | 8 |
| FOXP3 | SP97 | Rabbit | Thermo Scientific | 25-50 | 4-8 |
| GFAP | Polyclonal | Chicken | Abcam | 1500 | 4 |
| Granzyme B | EPR20129-217 | Rabbit | Abcam | 400-500 | 4 |

| | | | | | |
|---|---|---|---|---|---|
| HLA-DR | TAL-1B5 | Mouse | Santa Cruz | 200-600 | 4 |
| ICOS | EPR20560 | Rabbit | Abcam | 100-200 | 4 |
| IDO-1 | V1NC3IDO | Mouse | Invitrogen | 400-500 | 4 |
| Ki-67 | MIB-1 | Mouse | Dako | 50-150 | 4 |
| LAG-3 | 17B4 | Mouse | Novus Bio | 100-200 | 4 |
| MYL9 | EPR13012 (2) | Rabbit | Abcam | 800 | 8 |
| NaKATPase | H-3 | Mouse | Santa Cruz | 400 | 4 |
| PD1 | EPR4877(2) | Rabbit | Abcam | 200-500 | 4-8 |
| PD-L1 | IHC411 | Rabbit | GenomeMe | 150 | 4 |
| PD-L1 | 73-10 | Rabbit | Abcam | 500 | 4 |
| Podoplanin | D2-40 | Mouse | Agilent | 40 | 8 |
| S100 | 4C4.9 | Mouse | Thermo Scientific | 100-400 | 4 |
| Tryptase | AA1 | Mouse | Biolegend | 1600 | 4 |
| Vimentin | SP20 | Rabbit | Cell Marque | 100-200 | 4 |
| Vimentin | SP20 | Rabbit | Abcam | 250 | 4 |
| VISTA (B7-H5) | D1L2G | Rabbit | Cell Signaling | 200 | 4 |
